# Spinal Cord Elongation Enables Proportional Regulation of the Zebrafish Posterior Body

**DOI:** 10.1101/2024.04.02.587732

**Authors:** Dillan Saunders, Carlos Camacho, Benjamin Steventon

**Affiliations:** Department of Genetics, University of Cambridge, Cambridge, UK, CB2 3EH

## Abstract

Early embryos display a remarkable ability to regulate the patterning of tissues in response to changes in tissue size. However, it is not clear whether this ability continues into post-gastrulation stages upon cell commitment to distinct germ layers. Here, we performed targeted removal of neural fated cells in the zebrafish tailbud using multi-photon ablation. This led to a proportional reduction in the length of both the spinal cord and paraxial mesoderm in the tail, revealing a capacity to regulate tissue morphogenesis across multiple tissues to build a well-proportioned posterior body. Following analysis of cell proliferation, gene expression, signalling and cell movements, we found no evidence of cell fate switching from mesoderm to neural fate to compensate for neural progenitor loss. Furthermore, we found that tail paraxial mesoderm length is not reduced upon direct removal of an equivalent number of mesoderm progenitors, ruling out the hypothesis that neuromesodermal competent cells enable proportional regulation. Instead, reduction in the numbers of cells across the spinal cord reduces both spinal cord and paraxial mesoderm length. We conclude that spinal cord elongation is a driver of paraxial mesoderm elongation in the zebrafish posterior body and that this can explain proportional regulation of both tissues upon neural progenitor reduction.

## Introduction

Cells must coordinate their morphogenesis, differentiation, and growth to form embryonic tissues. In turn, tissue expansion must be coordinated such that each one forms with the correct proportions for the embryo. Investigations into the control of tissue proportions have predominantly focused on the specification of their primordia from a field of competent cells. Many of these patterning signals show a remarkable ability to scale with changes to the size of the field and have been thoroughly reviewed elsewhere (Čapek & Müller, 2019; Thompson et al., 2018). Tissue proportions can also be coordinated after their specification through the coupling of the morphogenesis of a tissue with that of its neighbours. Though less studied, this type of proportional regulation has been recently described in the elongation of the body axis in zebrafish (McLaren & Steventon, 2021; Tlili et al., 2019) and avian (Xiong et al., 2020) embryos. This phenomenon is an example of multi-tissue tectonics, in which the deformation of a tissue at the mesoscopic scale, known as tissue tectonics (Blanchard et al., 2009), can impact the dynamics of morphogenesis in neighbouring tissues (Busby & Steventon, 2021).

The ability of an embryo to regulate the proportions of its tissues when reduced in size is most evident prior to, and during gastrulation, and has been demonstrated in zebrafish (Almuedo-Castillo et al., 2018; Huang & Umulis, 2019; Ishimatsu et al., 2018), *Xenopus* (De Robertis, 2009), chick (Spratt Jr. & Haas, 1960) and mouse (Nichols et al., 2022). In the case of fish and frog embryos the resulting correctly patterned body plan does not recover its wildtype size, while some species such as mouse display both proportional and absolute size regulation (Nichols et al., 2022).

The formation and morphogenesis of the primary embryonic body axes occurs during gastrulation and then continues through a process known as posterior body elongation, which generates the embryo’s tail and a species-specific amount of its trunk (Steventon & Martinez Arias, 2017). The elongation of the tail tissues, such as spinal cord and paraxial mesoderm (somites and pre-somitic mesoderm), occurs through a combination of progenitor addition and subsequent morphogenesis of the tail tissues (Bénazéraf, 2019; Steventon et al., 2016). The raw materials for tissue extension are the progenitor cells located most caudally in the embryo in a structure known as the tailbud. There is now considerable evidence that some of these progenitor cells, often known as neuro-mesodermal progenitors (NMPs), are not restricted to either neural or mesodermal identity and can give rise to descendants that end up in either tissue type. The details of this have been reviewed extensively elsewhere (Wymeersch et al., 2021).

In zebrafish, the manipulation of Wnt signalling transduction in single cells determines whether they become neural or mesodermal (Martin & Kimelman, 2012) and from this we can infer that neuro-mesodermal competent cells are present in the tailbud. However, cell tracking at various developmental stages has found few or no progenitors producing both neural and mesodermal descendants (Attardi et al., 2018; Kanki & Ho, 1997; Lange et al., 2023). This is because NMP behaviour is dependent on the environment of the cells. For example, the low levels of cell division in the zebrafish tailbud (Attardi et al., 2018; Bouldin et al., 2014; Kanki & Ho, 1997; Steventon et al., 2016) mean that cells have few progeny to contribute to either tissue type. This has been proposed to account for differences in NMP behaviour between mouse and zebrafish (Sambasivan & Steventon, 2021; Wymeersch et al., 2021). Importantly, another factor that could affect NMP behaviour is the morphogenetic flow of the progenitor cells, which was first proposed to affect the prevalence of NMP cells in the early chick embryo as these cells are located at the divergence of two flows (Wood et al., 2019). Notably a similar difference in morphogenetic behaviour appears to separate dorsal and ventral located progenitors in the zebrafish tailbud (Lange et al., 2023; Lawton et al., 2013). This has led to the proposal to use the broader term neuro-mesodermal competent cells (NMC cells) when referring to these cells (Binagui-Casas et al., 2021).

Given that this competent cell population is retained in the zebrafish tailbud without it being utilised as a stem cell pool, it has been suggested that this population may enable zebrafish embryos to be robust to imbalance of neural and mesoderm progenitor allocation during gastrulation (Sambasivan & Steventon, 2021). In line with this hypothesis, the inhibition of Wnt signalling throughout the zebrafish embryo during this period leads to the loss of paraxial mesoderm but not the loss of neural tissue (Martin & Kimelman, 2012). However, there is a growing body of evidence that zebrafish posterior body elongation is driven primarily by the morphogenesis and growth of the tail tissues, anterior to the tailbud (McLaren & Steventon, 2021; Özelçi et al., 2022; Steventon et al., 2016; Tlili et al., 2019). Furthermore, studies have shown that these tissues are coupled together through the extra- cellular matrix (Guillon et al., 2020; Tlili et al., 2019). This raises the alternative hypothesis that multi-tissue tectonics could ensure the proportional elongation of neural and paraxial mesodermal tissue.

Consequently, we set out to investigate whether there is any capacity for the proportional regulation of neural and mesodermal tissue elongation in the zebrafish tail, and if so to determine whether this regulation is coordinated by changes in NMC cell behaviour or multi-tissue tectonics.

## Results

### 1. Loss of neural progenitors results in a proportional reduction in tail elongation

To investigate whether there is a capacity for regulation in the proportion of neural and mesodermal tissue in the tail we needed a method to reduce the number of cells in the system. We chose to use two-photon ablation as it allows the targeting of a subpopulation of cells in three-dimensions and acts rapidly to cause cell death (Lanvin et al., 2015; Vogel et al., 2005). Additionally, two-photon ablation has recently been utilised to investigate the role of notochord growth in zebrafish tail development (McLaren & Steventon, 2021).

We first set out to ablate the neural-fated progenitors in the tailbud at the 14 somite stage, to see if this affected the proportional formation of spinal cord relative to paraxial mesoderm. Neural-fated progenitors are located in the dorsal and posterior region of the zebrafish tailbud according to previous lineage tracing and fate-mapping studies (Attardi et al., 2018; Kanki & Ho, 1997; Lange et al., 2023; Lawton et al., 2013). We utilised these morphological criteria to localise the region of interest (ROI) for the ablations which is clearly within the *sox2* positive region of the tailbud and therefore includes the neural-fated progenitors (Fig. 1A; 2A). Mapping ablations together shows that there is a high precision in user selected ablation ROIs (Fig. 1B) meaning that we are reliably targeting neural progenitor cells.

**Figure 1.**
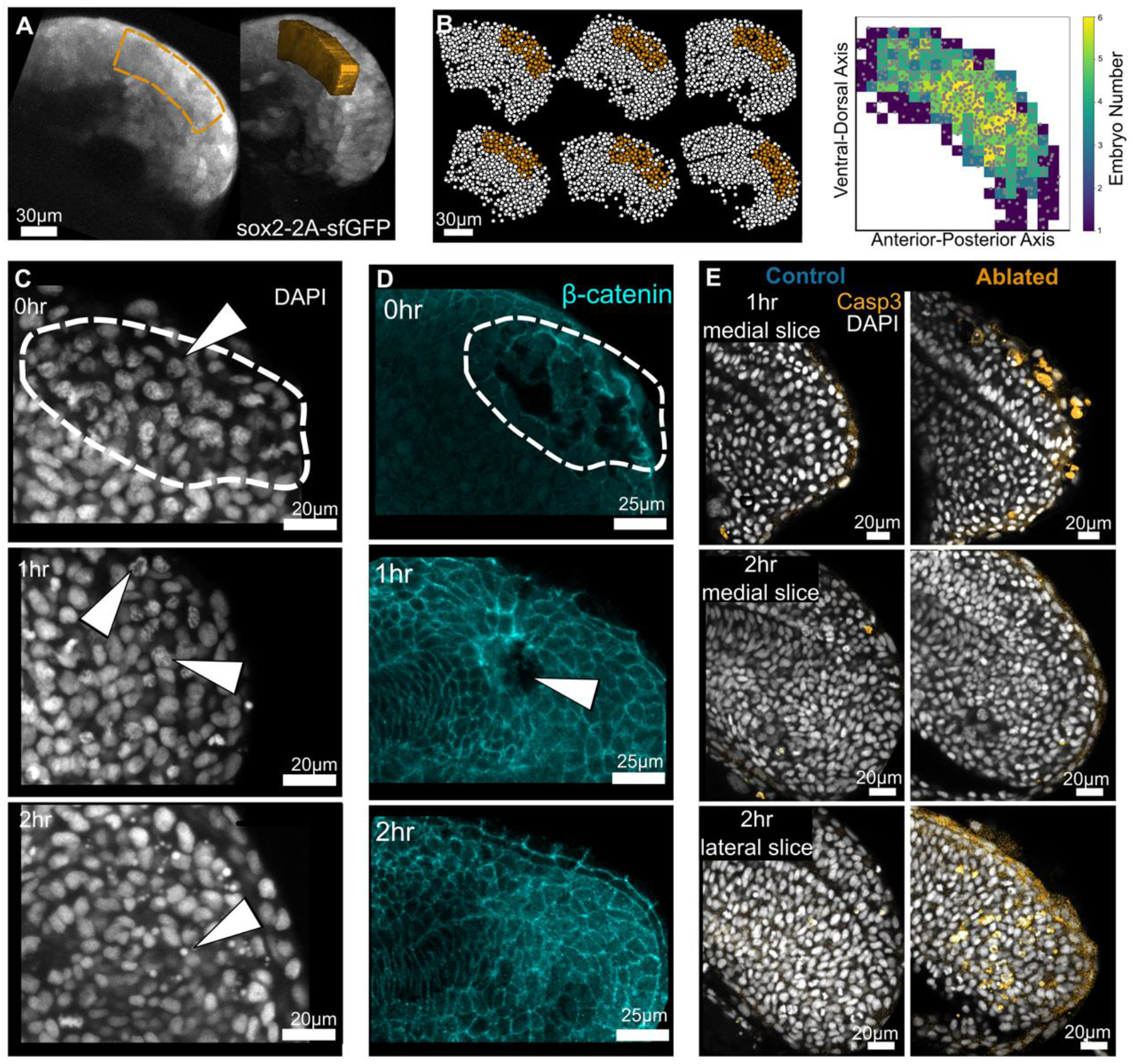
two-photon ablation causes localised cell death in neural-fated progenitor region. (A) Lateral and oblique views of a typical 3D ablation region of interest located in the neural fated progenitor region as indicated by the expression of the Sox2:GFP knock-in line. (B) Result of segmentation and registration of tailbud nuclei, ablated region in orange. The precision of the ablation location between embryos is displayed in the heatmap (n=6). (C) Representative images of DAPI stained nuclei fixed at successive timepoints post ablation. Nuclei in the ablated region (dashed line), become progressively more irregular and then condense as they undergo pyknosis (arrows). (D) Representative images of embryos fixed at successive timepoints post ablation with cell membranes marked by β-catenin. Cell membrane integrity is initially disrupted in the ablated region and gradually heals over time. 0hr: ablated, n=3; control, n=4. 1hr: ablated, n=5; control, n=5. 2hr: ablated, n=7; control, n=6. (E) Apoptotic cells marked by activated Caspase3 are seen after 1hr in the region of ablation. By 2hrs post ablation remaining apoptotic cells and debris are localised laterally as dead cells move out of the tailbud. 1hr: ablated, n=4; control, n=4. 2hr: ablated, n=6; control, n=6.

Next, to validate that two-photon ablation causes sufficient and rapid cell death under our experimental conditions we examined fixed samples at intervals following ablation. In these samples we observe irregular nuclear staining (Fig. 1C) and the destruction of cell membranes (Fig. 1D) immediately following ablation. This progresses to clear nuclear pyknosis (Fig. 1D) at one- to two-hours post ablation as the ablation decreases in size (Fig. 1D). At this stage staining for activated Caspase3 shows that these cells are undergoing apoptosis (Fig. 1E).

We then ablated neural-fated progenitors in groups of embryos using ROIs of increasing size, from which we quantified the number of nuclei in the ablated region as an estimate of the number of cells removed from the tailbud (Fig. 2B). These embryos were left to grow, alongside unablated control embryos, until the end of somitogenesis when they were fixed and stained for nuclei and actin to allow morphological identification of all tail tissues.

**Figure 2.**
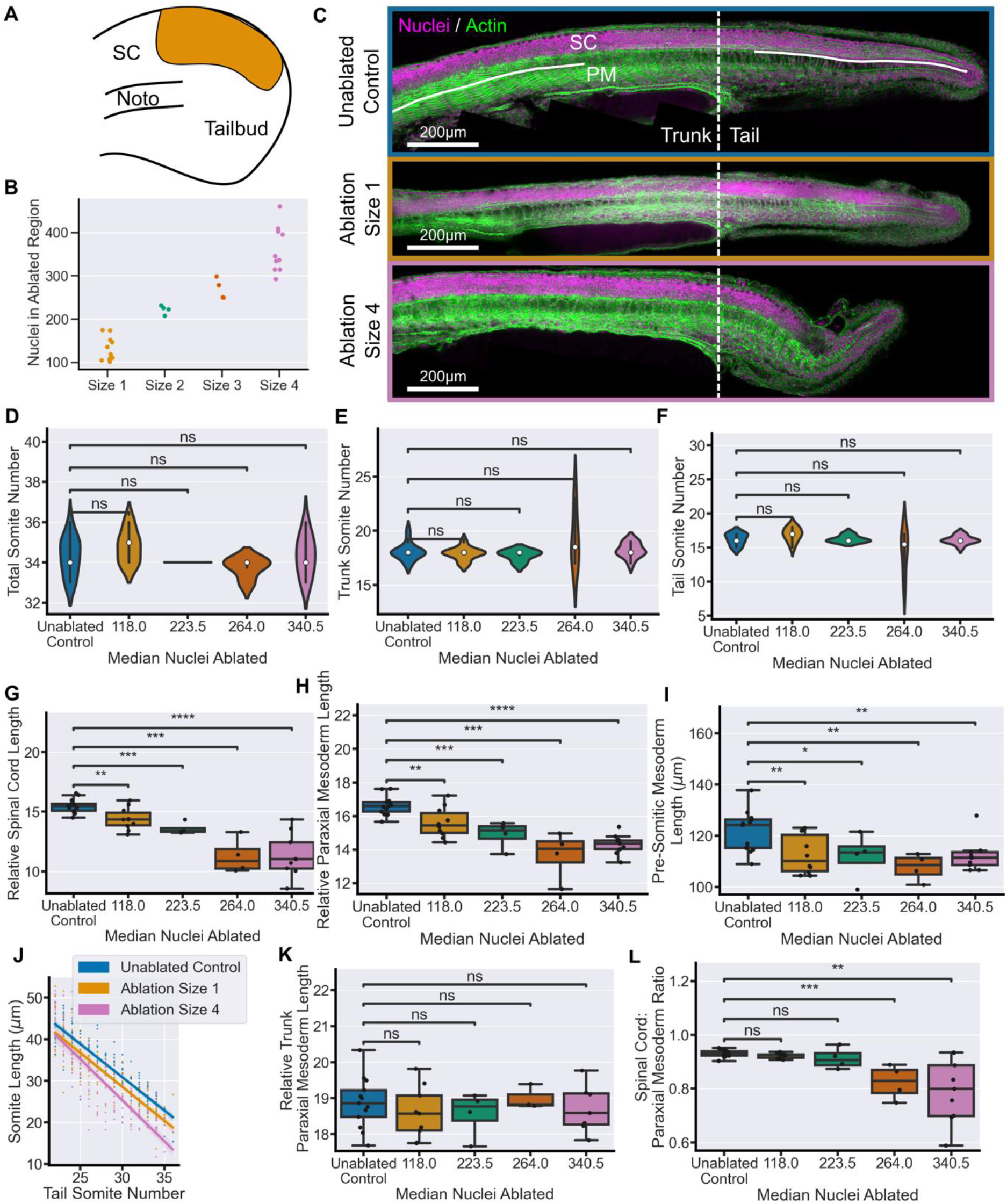
Neural-fated progenitor ablation results in a proportional reduction of tail tissue elongation. (A) Schematic showing an example neural-fated progenitor ablation in the 14-somite stage tailbud. (B) Number of nuclei in the ablation ROI prior to ablation with increased ROI size. (C) Representative examples of embryos at 30hpf stained for nuclei and actin. The morphology of the tail is comparable between size 1 ablations and unablated controls. While size 4 ablations cause a clear defect in tail formation. Solid lines indicate regions measured in (G) and (K). (D) Total somite counts, as well as (E) trunk, (F) tail somite number, are comparable between control embryos and all ablation conditions. (G) Spinal cord length, and (H) paraxial mesoderm length, measured from 22^nd^ somite, relative to total somite number, both show a significant decrease in all ablated conditions compared to controls. (I) Pre-somitic mesoderm length shows a significant decrease in all ablated conditions compared to controls. (J) Average somite length is consistently decreased in Size 1 ablation compared to controls. In Size 4 ablations there is a more notable decrease in the most posterior somites. (K) Trunk paraxial mesoderm length (2^nd^ to 10^th^ somites), relative to total somite number, shows no significant difference between any ablated condition and controls. (L) Spinal cord length relative to mesoderm length from 22^nd^ somite shows that ablations of size 1 and 2 maintain a ratio of tail tissues comparable to control embryos while size 3 and size 4 embryos have a significantly lower ratio. Unablated control, n = 13; size 1, n=10; size 2, n=4; size 3, n=4; size 4, n=9. Conditions were compared using Mann-Whitney-Wilcoxon test. *, p <=0.05; **, p <= 0.01; ***, p <= 0.001; **** p <= 0.0001. SC = spinal cord, Noto = notochord, PM = paraxial mesoderm.

Qualitatively, ablations of size 1 (118 nuclei) and 2 (223 nuclei) have little effect on the morphology of any of the tail tissues. We estimate, based on quantification of the number of *sox2* nuclei in the tailbud at this stage that these ablations account for 9-17% of *sox2* positive cells. Ablation of a larger proportion neural progenitors, size 3 (264 nuclei) and 4 (340 nuclei) (Fig. 2B), do cause a reduction in tail spinal cord size and overall tail elongation (Fig. 2C). We quantified the number of somites to determine whether ablation affects their production and found that, with one exception in the size 3 ablations, no ablation condition alters the total number of somites (Fig. 2D), or the number within either the trunk or the tail (Fig. 2E, F), as measured from the yolk extension. From this we can conclude that ablations do not affect the somitogenesis clock.

To quantify whether ablation affects the elongation of the spinal cord and/or paraxial mesoderm we measured the length of both tissues in the tail, beginning at the 22^nd^ somite which is always located in the tail. The length of each tissue was normalised to the number of somites to account for slight variation in developmental timing amongst embryos. We found that even the smallest ablations cause a significant decrease in the elongation of the tail spinal cord, with the effect even more pronounced following the largest ablations (Fig. 2G). This demonstrates that neural-fated progenitors in the tailbud are required for spinal cord elongation in the posterior body. Importantly, the loss of neural-fated progenitors also significantly effects the elongation of the tail paraxial mesoderm as well (Fig. 2H). This decrease in elongation is consistent in the pre-somitic mesoderm (Fig. 2I) as well as in the somites across the tail (Fig. 2J) but cannot be seen in the length of trunk somites which are comparable between conditions (Fig. 2K).

Comparison of the lengths of the two tissues relative to one another shows that the spinal cord and paraxial mesoderm lengths scale their elongation following ablations of size 1 (118 nuclei) and size 2 (223 nuclei). This proportional reduction breaks down following larger ablations, around 264 nuclei or more, as the elongation of the spinal cord is more effected by the large-scale loss of neural progenitors than the paraxial mesoderm (Fig. 2L). Taken together this demonstrates that the proportional extension of neural and mesodermal tissue in the tail has a considerable capacity for regulation following the loss of neural progenitors.

### 2. Sox2 and Tbxta expression is robust to loss of neural progenitors

NMC cells are a potential candidate for facilitating the regulation of neural and mesodermal proportions as they could balance the relative number of neural versus mesodermal fated progenitors. To investigate whether this is the case we fixed embryos at discrete intervals following size 1 ablations (on average 118 neural-fated nuclei) and stained them for the expression of the transcription factors *sox2* and *tbxta* (Fig. 3A). These genes are involved in the early steps of neural (Okuda et al., 2010) and mesodermal (Martin & Kimelman, 2008, 2010) differentiation respectively, and their co-expression is commonly associated with a cell in an NMC state (Binagui-Casas et al., 2021; Wymeersch et al., 2021). In zebrafish the *sox2/tbxta* positive cells that correspond to the NMC progenitors are located in the posterior wall of the tailbud (Attardi et al., 2018; Martin & Kimelman, 2012; Toh et al., 2022).

**Figure 3.**
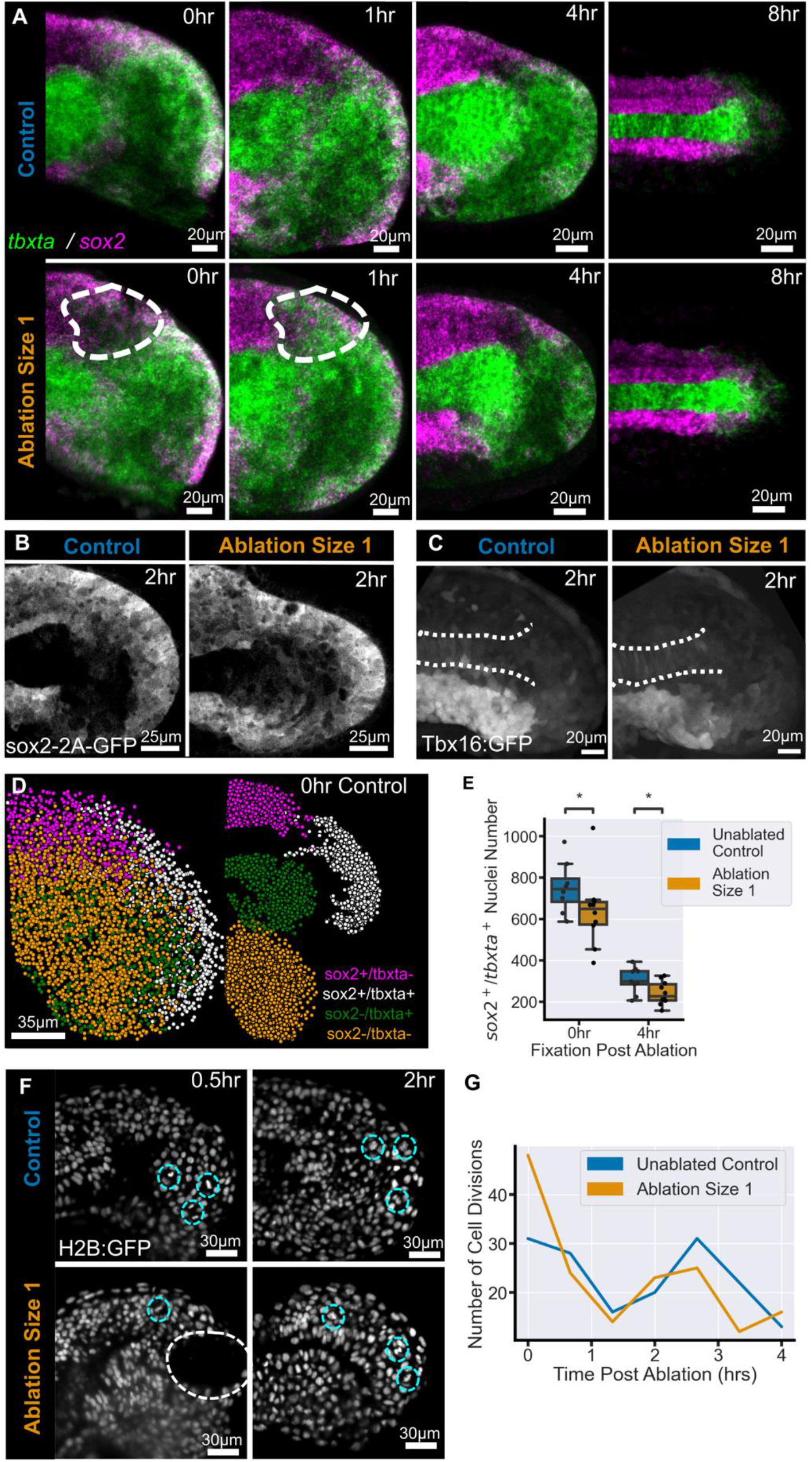
Neural progenitor ablation does not affect the gene expression pattern or cell division levels in the tailbud. (A) Mean (average) projection through the midline of representative images of the *sox2/tbxta* expression pattern in control and ablated embryos over time. Initial disruption of the pattern can be seen in ablated embryos at 0hr post ablation (white outline). By 4hrs, and up to 8hrs, post ablation the gene expression pattern is comparable to control embryos. 0hr: ablated, n=10; control, n= 8. 1hr: ablated, n=11, control, n=9. 4hrs: ablated, n=10; control, n=9. 8hr: ablated, n=3; control, n=3. (B) Sox2-GFP transgene expression visualised with GFP antibody shows expression in the posterior wall and spinal cord 2hrs after ablation, similar to controls. Ablated, n=7; control, n=7 (C) Tbx16-GFP transgene expression visualised with GFP antibody shows a comparable pattern between control and ablated embryos. Ablated, n=5, control, n=5. (D) Centroids of whole tailbud 3D nuclear segmentation of a 0hr control embryo using the mean expression of *sox2* and *tbxta* different cell populations within the tailbud can be isolated. (E) Number of *sox2/tbxta* double positive nuclei in each tailbud. Ablation causes a significant drop in the number of nuclei at 0hr post ablation compared to controls. A significant reduction remains at 4hrs post ablation compared to controls. (F) Long-term lightsheet imaging of an ablated embryo (white circle) and a stage-matched control allows manual identification of dividing cells (cyan circle), (H) there is no clear difference between the number of divisions in ablated and control embryos over 4 hours. Ablated, n=1; control, n=1. Conditions were compared using Mann-Whitney-Wilcoxon test. *, p <=0.05.

The time course of *sox2* and *tbxta* expression following ablation shows that there is an initial decrease in *sox2* expression in the location of the ablation immediately after it has occurred. One-hour post ablation some disorganisation of the gene expression pattern remains but by four hours post ablation ablated embryos are qualitatively similar to control embryos. Importantly, *sox2/tbxta* co-expressing nuclei are still located in the posterior wall of the tailbud. At the end of somitogenesis the *sox2/tbxta* cells have differentiated and a *sox2* positive spinal cord can be seen in both control and ablated embryos, while the tailbud remnants express *tbxta* (Fig. 3A). This suggests that the gene expression pattern in the tailbud is remarkably robust to the loss of neural progenitor cells.

To investigate this further we performed ablations in Sox2::GFP and Tbx16::GFP transgenic embryos, then fixed and stained the embryos after 2hrs for GFP to detect the activation of the transgenes. At this stage we observe that the ablation has little impact on the distribution of Sox2::GFP signal which remains strong throughout the posterior wall (Fig. 3B). Similarly, the expression of Tbx16::GFP remains localised to the PSM and the endothelial mesoderm with no evidence of any shift into the neural-fated region caused by the ablation (Fig. 3C).

Taken together these results show that neural progenitor ablation does not appear to significantly affect the gene expression pattern within the tailbud which suggests that NMC cells may not be regulating tissue proportioning.

### 3. NMC cells do not increase division levels in response to ablation

To be sure of whether NMC cells play a role in elongation regulation, we needed to quantify NMC cell behaviour in response to ablation. NMC cells could change their behaviour and balance out the progenitor population in two ways, either through increased levels of division or through a change to the balance of neural versus mesodermal differentiation.

Firstly, to determine whether NMC cells replenish the cells lost to ablation we investigated the *sox2-tbxta* pattern in greater detail by segmenting individual nuclei within the tailbud in three-dimensions and quantifying the mean intensity of each gene within each nucleus. Each nucleus can be represented as a single point and coloured according to whether it expresses *sox2* and/or *tbxta* (Fig. 3D).

We then quantified the number of *sox2/tbxta* cells in each tailbud at each timepoint to determine whether there is an increase associated with higher division levels. Instead we found that immediately following ablation there is a significant drop in the number of NMC cells, as expected from the loss of gene expression in the dorsal-posterior tailbud wall (Fig. 3A), however, this decrease is not recovered over time and remains significant after four hours post ablation (Fig. 3E). To further verify a lack of cell division we quantified the number of divisions occurring in the tailbud over the course of four hours following ablation (Fig. 3F) and found no difference compared to the number of divisions in the unablated tailbud (Fig. 3G). This suggests that neural progenitor ablation does not trigger increased cell division to replace the lost cells in the tailbud.

### 4. NMC cells do not shift their fate following ablation

Alternatively, NMC cells could be changing their position in the tailbud to repopulate the neural progenitors at the expense of mesodermal differentiation. In the zebrafish tailbud the localisation of the NMC cells is tightly linked to their pattern of differentiation with the dorsally located cells becoming neural and the ventrally located cells becoming mesodermal (Attardi et al., 2018; Toh et al., 2022). Therefore, in order to balance the loss of neural progenitor cells we would expect to see a shift from the mesodermal fated region towards the neural fated region, potentially as the result of cell rearrangements during ablation healing.

To explore whether this process occurs we live imaged embryos following ablation. Observation of LifeAct::GFP and MyosinII::mCherry transgenic lines allows visualisation of the ablation healing process occurring over one and a half hours (Fig. 4A, B). Of particular importance are the cells on the edge of ablation that move into the ablated region and form new cell-cell contacts (Fig. 4A; box). This healing process is also associated with increased activity of both Actin and non-muscle MyosinII at the edge of the cells exposed to the ablation. These features are suggestive of changes to local cell behaviour caused by ablation.

**Figure 4.**
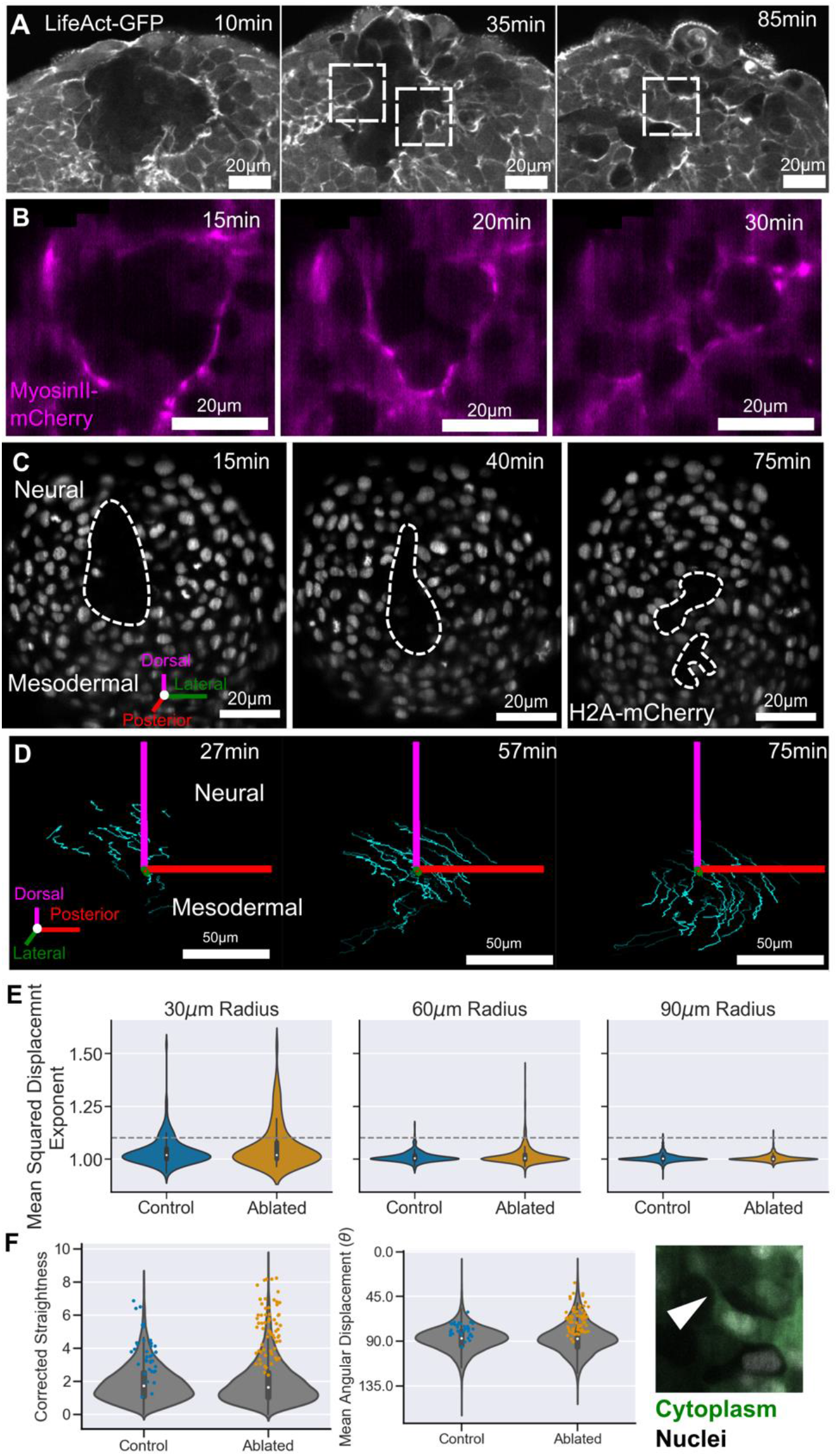
Ablation healing involves local increases in acto-myosin activity and more directional movement. Representative images of ablation healing visualised using (A) Lifeact:GFP, (B) MyosinII:mCherry, cells increase (A) actin, (B) myosin, levels at the ablation edge and move into the ablated region to re-establish cell contacts (boxes in (A)). (C) Reprehensive images of ablation healing using H2A:mCherry to mark nuclei, these nuclei were identified and tracked over one hour. Ablated, n=4; control, n=4. (D) a subset of tracks from one ablated embryo. (E) The exponent of the mean squared displacement for each track gives a measure of the consistency of motion. Tracks are grouped according to their starting distance relative to the ablation centre or equivalent control point. There are a greater number of tracks with a more consistent motion in ablated embryos close to the ablation compared with control embryos (above the grey-dashed line). These tracks (coloured points) also rank highly, compared to all tracks (grey), in other measures of directional motion, (F) corrected track straightness, (G) mean angular displacement. (H) Some directional tracks are associated with cells moving into the ablated region (arrow indicates cell protrusion crossing the ablated region).

To quantify this in greater detail we live imaged nuclei (Fig. 4C) and extracted tracking information describing their individual movements over time, in three dimensions (Fig. 4D). We then selected only those tracks which begin in the first 10 minutes of the movie and calculated their distance from the centre of the ablation or a point in an equivalent location of an unablated control embryo. A common metric for quantifying cell motion on a spectrum of diffusive to directional is the mean squared displacement (MSD). The exponent of the MSD curve gives the degree of directional motion at value greater than 1.0 (Beltman et al., 2009; Hu et al., 2023; Lawton et al., 2013). Plotting the distribution of this value for all tracks at different distances from the ablation centre (or equivalent) shows that the majority of tracks have an MSD exponent of around 1.0 which is indicative of random migration (Fig. 4E). A user defined threshold was set at an exponent of 1.1 to select tracks which have less random, and therefore more directional motion (grey dashed line). We observe that both ablated and control embryos contain a subset of tracks that have a relatively high MSD exponent. Importantly, there is a greater proportion of these tracks in ablated embryos and particularly among tracks which begin close to the ablation (Fig. 4E). These tracks also exhibit other measures of directional motion such as a high corrected straightness (Fig. 4F), and a high mean angular displacement (Fig. 4G). An example of a nucleus which produces one of these directional tracks is shown in Figure 4H, here the cell is protruding into the ablated region and forming a new cell-cell contact (arrowhead). Taken together this suggests that ablation does cause a small, local, increase in the amount of directional motion.

Despite the fact that ablation healing does alter the behaviour of a subset of nuclei we noticed that, even as it is healing, the ablation is displaced towards the region of mesodermal fate (Fig. 4C). Similarly, the tracks, both those that display more directional movement (Fig. 4D), and those that don’t (Fig. 5A), all display a clear neural to mesodermal flow over the course of the movie. This would suggest that the ablation is not sufficient to cause a mesodermal to neural shift in the fate distribution in the tailbud. To confirm this, we calculated the percentage of tracks for each embryo within 60μm of the ablation that are displaced in either the neural or mesodermal direction. We find that the majority of tracks in each embryo are displaced towards the mesodermal fated region in both control and ablated embryos (Fig. 5B). Consequently, we would not expect there to be a change in the balance of neural versus mesodermal fated cells in the tailbud following ablation.

**Figure 5.**
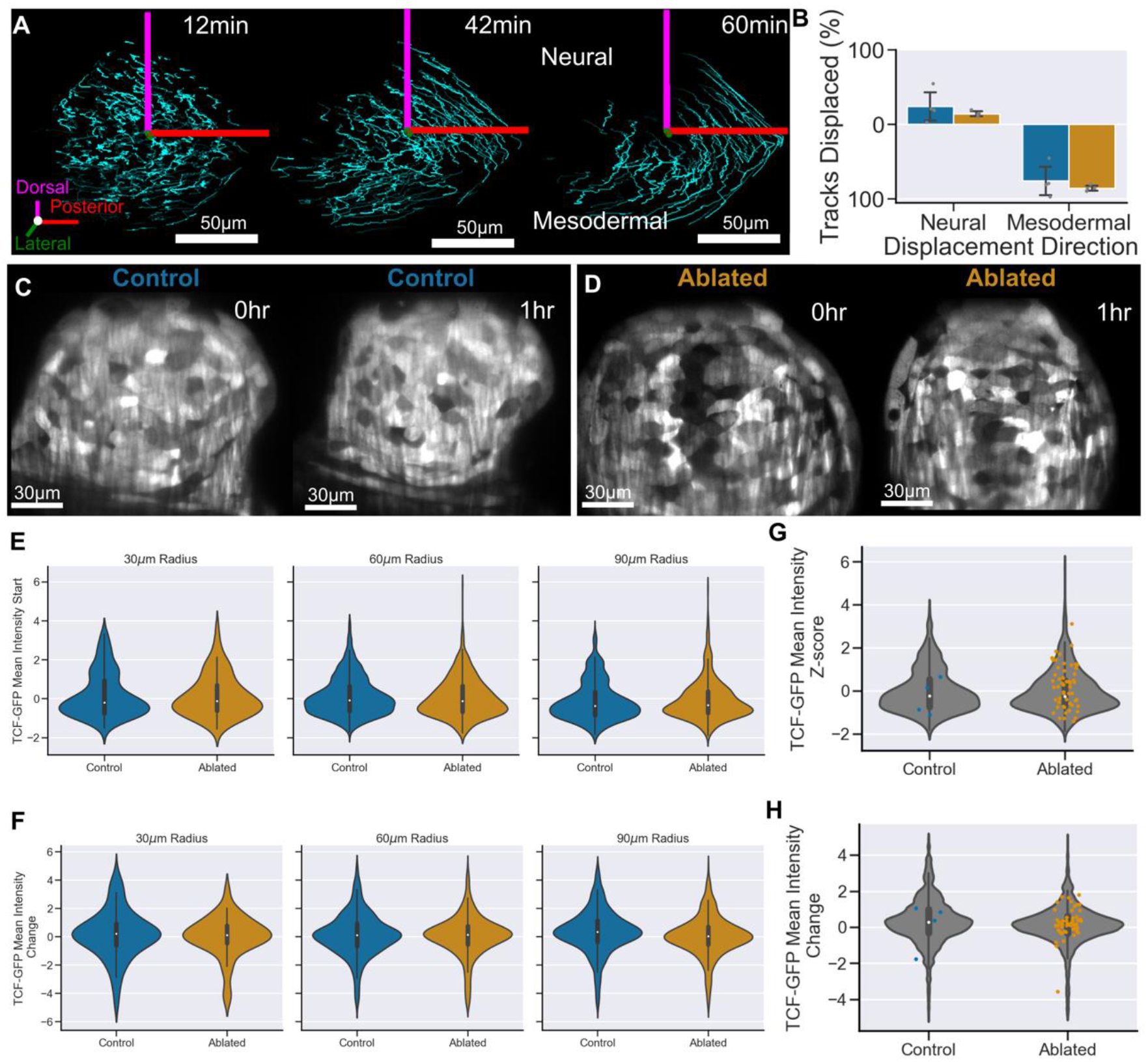
Global cell flow and Wnt signalling transduction are robust to neural progenitor ablation. (A) Representative tracks from an ablated embryo which start within 60μm of the ablation. (B) The vast majority of tracks are displaced in the mesodermal direction in both control and ablated embryos. Ablated, n=4; control, n=4. Representative images of TCF-GFP highlighting the level of Wnt signalling transduction in (C) control, and (D) ablated embryos. (E) TCF-GFP intensity, z-score normalised for each embryo, at the start of each track for tracks at different distances from the ablation/reference point. Tracks close to the ablation have comparable levels of TCF-GFP to control embryos. (F) Mean TCF-GFP intensity change over the track show both increase and decrease in Wnt transduction with ablated embryos having a similar distribution to control embryos. (G) TCF-GFP starting intensity, and (H) TCF-GFP mean intensity change, for all tracks within 60μm of the ablation (grey) overlaid with the intensity of the directional tracks (coloured) shows that directional tracks do not have a bias in Wnt transduction. Ablated, n=2; control, n=2.

Further indication that ablation does not trigger a global change in the balance of progenitors is in the lack of change to Wnt signalling in the tailbud following ablation. Wnt is the key signal in the neuro-mesodermal fate decision (Martin & Kimelman, 2012) so we imaged embryos with a TCF::GFP transgene to visualise the transduction of the signal. We observe that there is great variation in the level of Wnt transduction as reported by this transgene in both control and ablated embryos (Fig. 5C, D). This is confirmed by the broad distribution of normalised mean intensity values at the start of each track (Fig. 5E). We then calculated the mean change in intensity across the track lifetime and find that the vast majority of tracks do not change their level of TCF::GFP in either control or ablated embryos, with a minority of tracks in both conditions either increasing or decreasing their Wnt transduction (Fig. 5F). Additionally, more directional cell motion does not correlate with a particular level of Wnt signalling (Fig. 5G) or a significant change in the level of Wnt transduction (Fig. 5H).

Taken together, this demonstrates that cells which heal the ablated region are the neighbouring, neural-fated cells rather than more distantly located mesodermal-fated cells. Without a shift in cell localisation or Wnt signalling transduction it is unlikely that NMC cells would be able to alter their differentiation pattern. Therefore, we conclude that NMC cells do not alter their differentiation in response to ablation in a way that would facilitate the regulation of neural and mesodermal tissue proportions.

### 5. Paraxial mesoderm elongation can withstand the loss of a significant number of mesoderm progenitor cells

As a final challenge to the hypothesis that a re-distribution of mesoderm fated cells towards neural enables proportional regulation upon neural progenitor loss, we asked if mesoderm progenitor ablation results in the same shortening of the tail paraxial mesoderm as we observe in neural ablations (Fig. 2). We ablated paraxial mesoderm progenitors on one lateral side of the tailbud (Fig. 6A). As with neural progenitor ablations we performed ablations of different sizes – 1 to 3 (Fig. 6B) and grew the embryos until the end of somitogenesis (Fig. 6C). Similar to neural progenitor ablations there are no morphological defects at the initial ablation sizes 1 and 2. These ablation size count for 14% of the average number of *tbxta* positive nuclei in the tailbud at this stage. At larger ablation sizes several somites are lost from the tail on the side of ablation (Fig. 6C,D). Notably, counting somites on the unablated side shows that large mesodermal ablations only affect the somitogenesis on one side of the tail (Fig. 6D).

**Figure 6.**
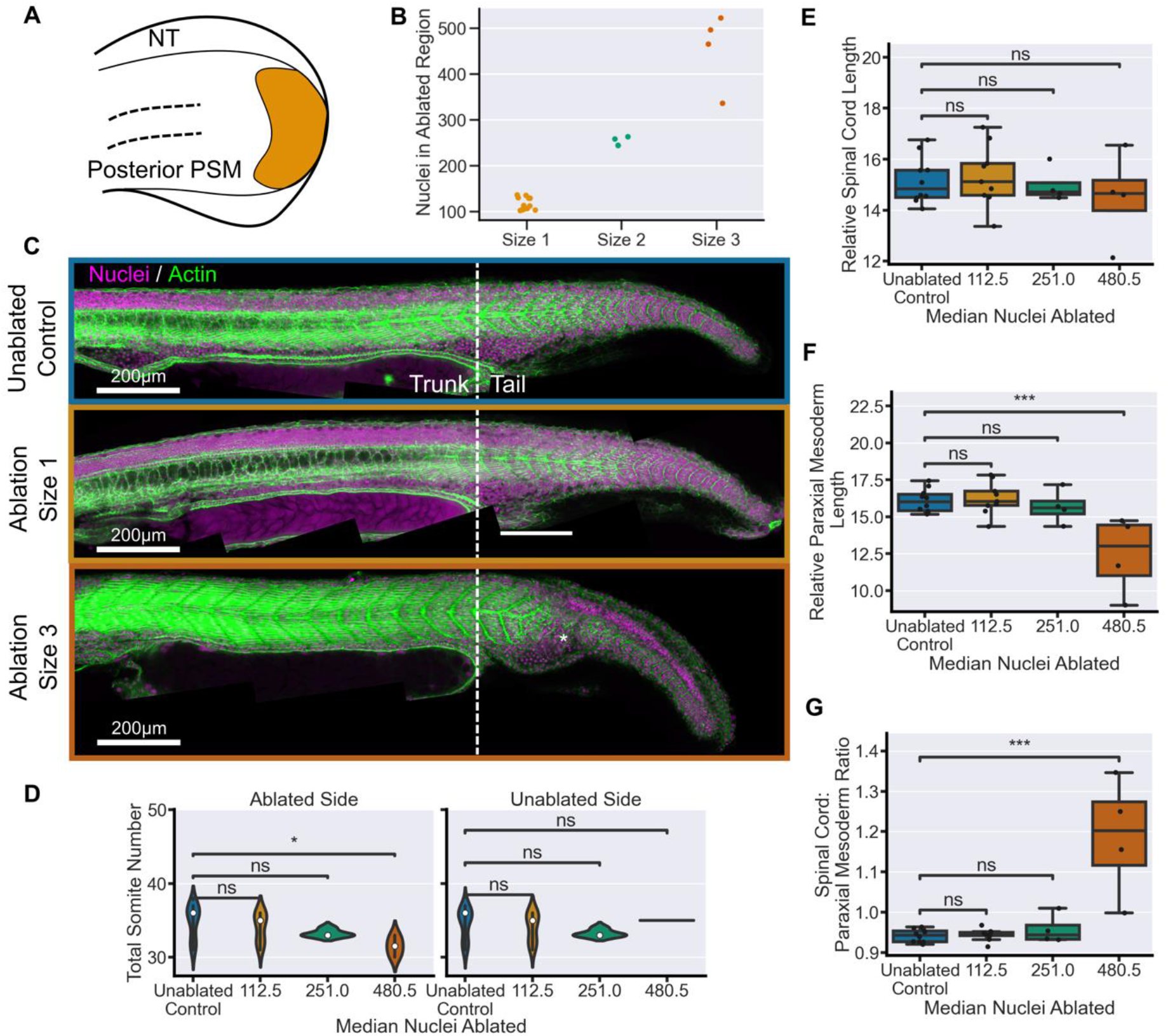
Mesoderm fated progenitor ablation does not affect tissue elongation. (A) Schematic showing the location of an example mesodermal fated progenitor ablation in the 14 somite stage tailbud. (B) Number of nuclei in the ablation ROI prior to ablation with increased ROI size. (C) Representative examples of embryos at 30hpf stained for nuclei and actin. The morphology of the tail is comparable between size 1 ablations and unablated controls. While size 3 ablations cause a clear defect in somite formation (asterisk). (D) Total somite count for both sides of the bilateral paraxial mesoderm. Size 1 and Size 2 ablations have comparable numbers of somites on both sides to control embryos. In size 3 ablations there is a loss of somites on the ablated side but the contra- lateral side remains unaffected compared to controls. (E) Spinal cord length, and (F) paraxial mesoderm length, measured from 22^nd^ somite, relative to total somite number, both show no significant decrease in size 1 and 2 ablations compared to controls. Size 3 ablations do have a significant difference in paraxial mesoderm length only. (G) Spinal cord length relative to mesoderm length from 22^nd^ somite shows that ablations of size 1 and 2 maintain a ratio of tail tissues comparable to control embryos while size 3 embryos have a significantly higher ratio. Control, n=10, size 1, n= 9; size 2, n=4; size 3, n=4. Conditions were compared using Mann-Whitney-Wilcoxon test. *, p <=0.05; **, p <= 0.01; ***, p <= 0.001. SC = spinal cord, PSM = pre-somitic mesoderm.

Importantly, in contrast to neural progenitor ablation, there is no reduction in the posterior body of spinal cord (Fig. 6E) or paraxial mesoderm (Fig. 6F) length following mesodermal progenitor ablations of up to 250 cells. A reduction in paraxial mesoderm elongation is only seen following the largest ablations of around 480 cells where it causes a deregulation of tissue proportions, as spinal cord elongation is not affected (Fig. 6G). This is particularly interesting as it shows that up to a certain threshold the elongation of the paraxial mesoderm is not dependant on the number of progenitors. Furthermore, a reduction in paraxial mesoderm elongation does not appear to affect the elongation of the spinal cord as significantly as the spinal cord effects the paraxial mesoderm. Therefore, the redistribution of mesoderm progenitors towards and neural fate cannot explain the proportional regulation in the tail that we observe in spinal cord ablations.

### 6. Genetic ablation of anterior spinal cord cells causes a reduction in neural and mesodermal tissue length

In the absence of changes to NMC cell behaviour following neural progenitor ablation we turned to investigate our alternative hypothesis, that spinal cord morphogenesis drives tail elongation and consequently tissue proportions. If this is the case, then a reduction in spinal cord morphogenesis anterior to the tailbud should also affect the length of the tail tissues. To reduce spinal cord elongation along the whole body we turned to a genetic ablation technique for tissue-specific expression of the bacterial Kid toxin (Labbaf et al., 2022). The expression of UAS:Kid in the spinal cord was driven by an Adamts3:GAL4FF (ATS3) transgene which has strong expression in motor neurons (Asakawa et al., 2013; Wang et al., 2020).

Following the crossing of the GAL4-UAS lines we observe a clear effect on the morphogenesis of the body axis (Fig. 7A) which is comparable to the effect of ATS3 mutants (Wang et al., 2020). Ablation of ATS3 positive cells causes a dorsal bending of the body axis which seems similar to the dorsal bending of the tail observed in large neural progenitor ablations (Fig. 2C). In the spinal cord, we observe pyknotic nuclei further confirming the action of the Kid toxin (Fig. 7B; arrows). Other tail tissues remain unaffected though the notochord is consistently kinked in the tail, which could indicate that its full elongation is being prevented (Fig. 7A; asterisk).

**Figure 7.**
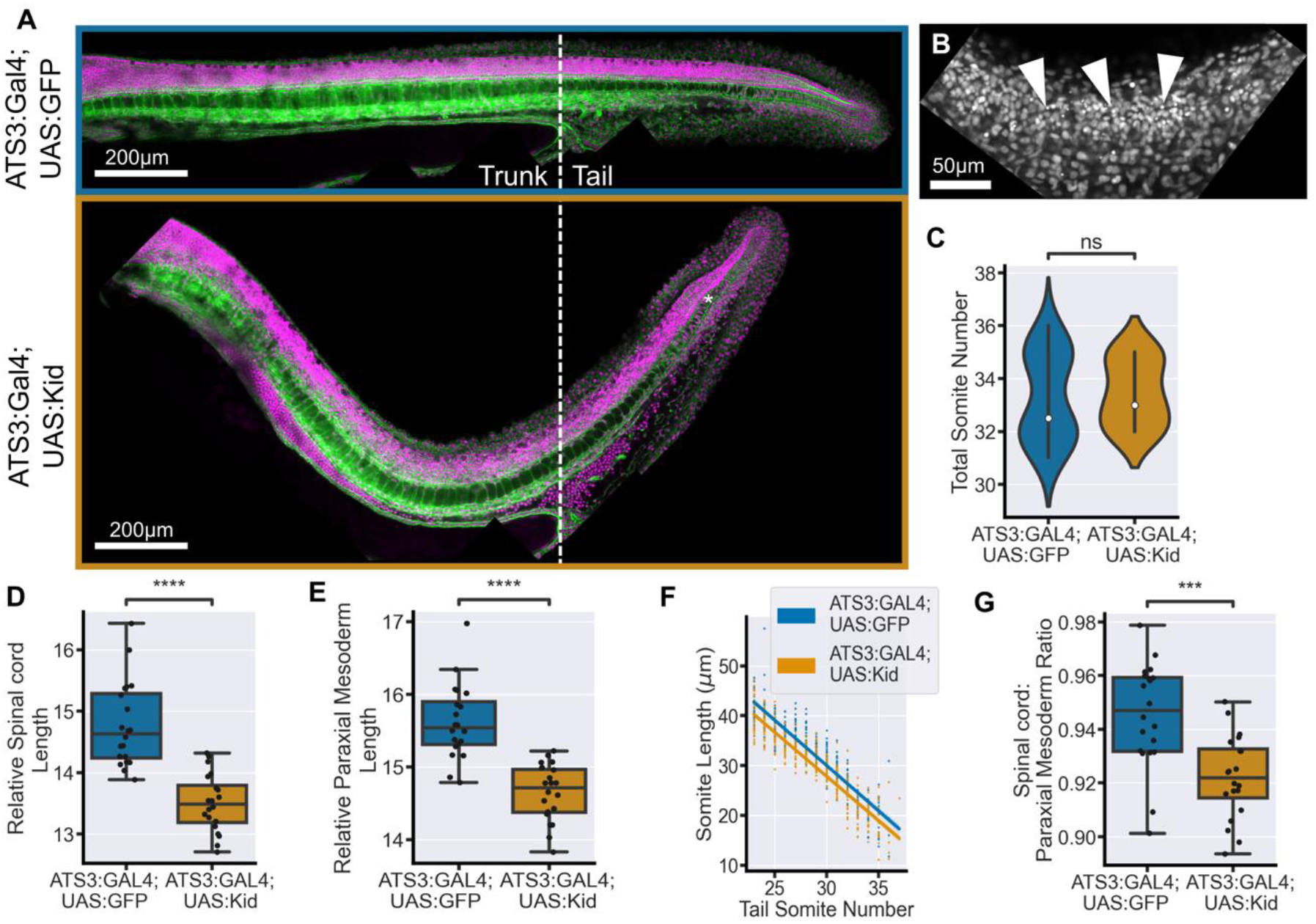
Genetic ablation of anterior spinal cord cells results in a proportional reduction in tail tissue elongation. (A) Representative examples of embryos at 30hpf stained for nuclei and actin. ATS3:Kid embryos have a notable dorsal bend in the body axis compared to controls. The notochord is often kinked in the tail (asterisk). (B) Pyknotic nuclei can be observed in the spinal cord particularly in the location of the axis bend. (C) Total somite counts shows no significant difference between ATS3:Kid and ATS3:GFP embryos. (D) Spinal cord length, and (E) paraxial mesoderm length, measured from 22^nd^ somite, relative to total somite number, both show a significant decrease in ATS3:Kid embryos compared to ATS3:GFP controls. (F) The decrease in paraxial mesoderm length is consistent across all the tail somites. (G) Spinal cord length relative to mesoderm length from 22^nd^ somite shows that spinal cord elongation is more affected than paraxial mesoderm elongation in ATS3:Kid embryos. ATS3:Kid, n=20; ATS3:GFP, n=20. Conditions were compared using Mann-Whitney-Wilcoxon test. *, p <=0.05; **, p <= 0.01; ***, p <= 0.001; ****, p <= 0.0001.

We do not see an effect on the number of somites in the tail following genetic ablation of spinal cord cells (Fig. 7C). However, we do observe a decrease in spinal cord length measured from the 22^nd^ somite in the tail in ablated embryos compared to controls (Fig. 7D). This demonstrates that the ablation of spinal cord cells effects the correct elongation of the tail spinal cord. We also find that there is a reduction in the length of the paraxial mesoderm in the tail (Fig. 7E), a decrease which is consistent across tail somites (Fig. 7F). This result demonstrates that elongation of the spinal cord secondarily affects the elongation of the paraxial mesoderm. By comparing the decrease in elongation of the spinal cord with the decrease in the elongation of the paraxial mesoderm we show that, similar to large two- photon ablations the spinal cord is more affected than the paraxial mesoderm (Fig. 7G).

## Discussion

In summary, our results show that there is a capacity for the regulation of tail elongation which ensures the proportional extension of the tail tissues. This demonstrates that the ability of the embryo to coordinate the scaling of tissue formation continues after pre-gastrulation stages where reduction in blastoderm size results in the complete scaling of all embryonic tissues (Almuedo-Castillo et al., 2018; Ishimatsu et al., 2018).

Notably, proportional regulation at pre-gastrulation stages occurs through changes in the pattern of cell differentiation through scaling of the signalling network (Almuedo-Castillo et al., 2018). In contrast, we demonstrate that the pattern of differentiation of the unspecified cells in the tailbud, the NMC cells, is not affected by the loss of neural fated progenitors. Neither is there a shift in progenitor morphogenesis or the transduction of the key patterning Wnt signal. Consequently, ablation results in only a reduction of the number of NMC cells in the tailbud. It is a notable result, that the progression of NMC cell differentiation is not dependent on the overall number of progenitors, which fits with previous work that NMC cells generate their own permissive signalling environment (Bouldin et al., 2015; Martin & Kimelman, 2008).

The lack of NMC cell response to ablation is in contrast to the loss of tissue proportions observed when manipulating Wnt signalling at tailbud stages (Martin & Kimelman, 2012). This raises an important distinction between what a cell can do and what a cell will do in the context of its environment. A similar distinction has been previously noted in relation to NMC cell behaviour and global division levels (Sambasivan & Steventon, 2021; Wymeersch et al., 2021). In this case the strong dorsal-ventral flow of cells in the tailbud (Lange et al., 2023; Lawton et al., 2013) is not sufficiently disturbed by the ablation and therefore does not result in a change in the contribution of NMC cells to neural versus mesodermal fate. A similar process of diverging cell flows has been proposed to split neural and mesodermal fated NMC cells in the early chick embryo (Wood et al., 2019).

Despite progenitor dynamics being highly robust to ablation, the correct elongation of the tail is sensitive specifically to the number of neural progenitors in the tailbud or the spinal cord. In contrast paraxial mesoderm elongation is not so easily affected by the loss of mesoderm progenitors. This result aligns with the known dynamics of tissue elongation in the tail in which the spinal cord undergoes the most volumetric growth, while the paraxial mesoderm elongates through ‘thinning and lengthening’ (Steventon et al., 2016). Consequently, loss of neural progenitors as the raw material for volumetric growth has an outsized effect on spinal cord formation. Reduced spinal cord elongation, in turn, affects the elongation of the paraxial mesoderm and thus maintains tissue proportions. This affect is likely transmitted through the mechanical coupling of the extra-cellular matrix as has recently been described in several works (Dray et al., 2013; Guillon et al., 2020; Tlili et al., 2019). Together, this positions spinal cord formation as a driver of posterior body elongation in zebrafish.

Finally, our results demonstrate the importance of considering the effect of multi-tissue tectonics (Busby & Steventon, 2021) on developmental processes. It is important to note that the exact interactions between the tissues is likely to vary across developmental time, space, and evolution so that the contribution of the morphogenesis of each tissue to posterior body elongation is not static or absolute. Notochord elongation may play a more prominent role at later stages (McLaren & Steventon, 2021), while in avian species for example, the morphogenesis of the PSM has an effect on the elongation of the spinal cord (Xiong et al., 2020). Overall, this work contributes to an expanding body of evidence that the formation and growth of the tail tissues, rather than the action of the progenitors themselves, are a major driver of posterior body elongation in zebrafish (Guillon et al., 2020; McLaren & Steventon, 2021; Özelçi et al., 2022; Tlili et al., 2019).

## Materials and Methods

### Animal Husbandry

The maintenance of adult zebrafish, including any regulated procedures, was conducted in accordance with the Animals (Scientific Procedures) Act 1986 Amendment Regulations 2012, following ethical review by the University of Cambridge Animal Welfare and Ethical Review Body (AWERB). Standard E3 media was used to culture embryos in all experiments except for those involving lightsheet microscopy in which methylene blue was omitted. Embryos were staged according to (Kimmel et al., 1995). To prevent involuntary embryonic muscle contraction tricaine (ethyl 3-aminobenzoate; Sigma #A-5040) was added at 0.16 g/ml in E3 medium. Transgenic zebrafish lines used: Tg(actb2:H2a-mCherry), Tg(actb2:H2b- GFP), Tg(actb2:Lifeact-EGFP) (Behrndt et al., 2012), Tg(sox2:2a-sfGFP) (Shin et al., 2014), Tg(β-actin:myl12.1-mCherry) (Maître et al., 2012), and Tg(7xTCF-Xla.Sia:GFP) (Moro et al., 2012).

### Microscopy

#### Two-photon microscopy

Embryos were mounted as in (Hirsinger & Steventon, 2017) . Embryos were imaged and ablated, using a Trim Scope II upright two-photon microscope (LaVision Biotec) with a 25x (1.05NA) water immersion lens. For ablation, the laser was set to 900nm and the imaging window was set to 200x200 in order to achieve a dwell time greater than 9μs. Laser power ranged from 1.3 to 0.5 Watts during the course of experiments, ablation was carried out with 80% laser power. The embryo was imaged for at least one complete stack prior to and after ablation. A successful ablation was determined by the presence of a small amount of recoil of adjacent nuclei in the timepoints following ablation.

#### Lattice Lightsheet microscopy

A V-shaped mould was created using 1% agarose in a 35mm glass-bottomed dish. Individual embryos were aspirated in 1% low melting point agarose and placed in the bottom of the V-shaped mould. An eye-lash tool was used to orient the embryo ventrally so that the tailbud was pressed against the bottom of the dish. Once the agarose set the embryo was covered with E3 medium and tricaine. The embryos were then imaged on a Lattice Lightsheet 7 (Zeiss) using inverted water immersion single illumination (13.3x; NA0.4) and detection (44.83x; NA1.0) objectives. A full stack was taken every 30 seconds at 28°C. Following imaging, embryos were removed from agarose and allowed to grow up at 28°C in E3 media overnight to confirm that mounting does not have an adverse effect on embryo growth or morphology.

#### Confocal microscopy

Fixed embryos were dissected completely away from the yolk and the head removed to ensure they lay flat on their lateral axis. They were mounted in VectaShield (Vector Laboratories) mounting media between two 1H coverslips stuck together with double sided tape. All fixed embryos were imaged on an inverted Zeiss LSM700. For embryos at 30hpf the whole embryo was imaged in sections using a 20X objective, with 4x line averaging. Images of the tailbud were taken using a 40X objective, with 2x line averaging. Images were collected with the same laser intensities, gain, pixel resolution for each experiment.

### Fixing and Staining

Embryos were fixed in 4% paraformaldehyde (PFA) (for 1 hour on the bench or up to 24hrs at 4°C. Embryos were then washed into phosphate buffered saline without magnesium and calcium ion with 0.05% Tween (PBST (-/-)). Nuclei were stained with 1:500 DAPI and actin was stained with 1:1000 Phalloidin tagged with Alexa Fluor 647nm in PBST for at least 24hrs. Individual two-photon ablated embryos were stained separately to be able to link the ablation with the final morphology. To visualise mRNA or proteins in situ embryos were pooled into ablated and control groups following ablation. Fluorescent in situ hybridisation was done using the Hybridisation Chain Reaction method as described in (Choi et al., 2018). Immunohistochemistry was carried out according as described in (Sorrells et al., 2013). Primary antibodies used were chicken anti-GFP (1:200), rabbit anti-activated Caspase3 (1:500), and mouse anti-βCatenin (1:200). All primary antibodies were raised in goat. Secondary antibodies were used at a concentration of 1:1000. DAPI was added at the end of the protocol at 1:500.

### Genetic ablation

ATS3:GAL4 (Wang et al., 2020) were crossed with either UAS:Kid (Labbaf et al., 2022) or UAS:GFP fish to produce embryos that ATS3:GAL4;UAS:Kid and ATS3:GAL4;UAS:GFP. This causes the expression of the bacterial Kid toxin or GFP in AdamTS3 expressing cells of the spinal cord. Embryos were selected based on phenotype.

### Image Analysis

#### Visualisation and registering

Images were visualised using FIJI (Schindelin et al., 2012), or Napari (Sofroniew et al., 2022). Two-dimensional length measurements were made using FIJI’s segmented line ROI tool. For measurements of tail length, images were collected individually and stitched together using a bespoke script for successive rounds of pairwise stitching utilising the FIJI Stitching plugin (Preibisch et al., 2009). Where necessary images were rotated using the TransformJ plugin (Meijering et al., 2001). Registration of ablated tailbuds was performed using ZebReg as described in (Toh et al., 2022). Data was visualised using the Python 3.9 packages Matplotlib and Seaborn. Statistical analysis was performed using the packages Scipy.Stats and StatAnnotations. Boxplots are set to display the median and quartiles of the data. Lineplots show the mean of the data and the 95% confidence interval. The functions used in image analysis can be viewed on GitHub https://github.com/DillanSaunders/TailbudAnalysis.

### 3D Nuclear Segmentation

Prior to segmentation nuclei images were pre-processed. First intensities were equalised using adaptive histogram equalisation with a 3D kernel cube of length 10μm. Second the images were filtered using a difference of gaussian filter (low sigma =1, high sigma = 3). Segmentation was performed in two steps first the nuclei were segmented in 2D using the StarDist 2D default model (Schmidt et al., 2018). 2D labels were then joined in 3D using a threshold of 0.6 intersection over union using a function adapted from CellPose (Stringer et al., 2021). The HCR images of fluorescent mRNA were then pre-processed using two rounds of median filtering (kernel = 0.8μm^3^). The median intensity values for *sox2* in the notochord and *tbxta* in spinal cord were used as an estimation of background intensity for each channel. This was used to filter out nuclei with background levels of both genes. The nuclei of the hypochord which are *sox2/tbxta* positive were then manually removed by eye.

#### Cell tracking

Images acquired from the Zeiss lattice lightsheet are saved into a single .czi file. These files were opened in ZenBlue software, cropped to the correct size and re-saved as .czi files. As the tail is prevented from growing outwards during this time these images do not need global registration. The images were opened in FIJI and converted to HDF5 data format (with deflate compression) using the plugin BigDataViewer (Pietzsch et al., 2015). Following conversion to HDF5/XML format the lightsheet movies were opened in the FIJI plugins MaMut (Wolff et al., 2018) or Mastodon (Pietzsch et al., 2014/2023). MaMut was used to generate a sample of manual tracks to be able to accurately parameterise automatic tracking in Mastodon. Automatic spot detection used a diameter of 10μm with a quality threshold of 50. Automatic spot linking was performed using a displacement of 10μm and a frame gap of no more than 2 frames. Cell division was permitted with the distance set to 15μm. Metrics of cell motion are well established in the field of cell migration and were calculated as described in (Beltman et al., 2009; Hu et al., 2023; Lawton et al., 2013).

## Acknowledgments

We would like to thank Martin Lenz and Kevin O’Holleran for assistance with the microscopes, Erez Raz for sharing the UAS:Kid plasmid, and Qiyu Chen for valuable feedback.

## Author Contributions

Conceptualization, D.S and B.S; Methodology, D.S, C.C, and B.S.; Investigation, D.S. and C.C; Writing, D.S, C.C, and B.S.; Visualization, D.S; Funding Acquisition, D.S, and B.S.

